# A Versatile Tool for Precise Variant Calling in Mycobacterium tuberculosis Genetic Polymorphisms

**DOI:** 10.1101/2023.07.24.550283

**Authors:** Safina Abdul Razzak, Zahra Hasan, M. Kamran Azim, Akbar Kanji, Sadia Shakoor, Rumina Hasan

## Abstract

**Background:** Whole genome sequencing (WGS) facilitates the diagnosis of multidrug-resistant MDR-TB through the interpretation of sequence variations (SV) in *Mycobacterium tuberculosis* (MTB) genes. Information on phenotypic and genotypic resistance associations continues to evolve, it is important to identify SV within genes of interest. We developed an MTB-VCF variant calling pipeline that can compare against the reference genome for any gene of interest. We demonstrate its utility for calling SV in genes associated with Rifampicin (RIF), Isoniazid (INH), Ethambutol (EM), and Streptomycin (SM) resistance.

**Methods:** MTB-VCF is a Python-based command line Variant Calling pipeline designed to streamline batch processing from raw reads (FastQ) files. SV called by MTB-VCF were compared with those identified by TBProfiler, KVARQ, CASTB, Mykrobe Predictor and Phy-ResSE pipelines. The sensitivity and Specificity of MTB-VCF SV calling were calculated against the drug susceptibility testing (DST) phenotype.

**Results:** MTB-VCF identified 868 SV present in 200 phenotypically resistant MDR-TB isolates. These were across *rpsl, rrs, rpoB, inh*A, *kat*G, *ahp*C, *gid*B and *emb*CAB genes. Of these, 684 SV were known to be associated with a resistance genotype, leading to a specificity of 97.75%. The SV called by the MTB-VCF was compared separately to resistance genotypes called by TB-Profiler, KvarQ, CASTB, Mykrobe Predictor, and PhyRes-SE pipelines, demonstrating a sensitivity of 99.5%.

**Conclusion:** The MTB-VCF pipeline offers a rapid and accurate solution for identifying SV in target genes for interpretation later. It can be run in large batches, proving flexible computing that allows for the customization of core bioinformatic pipelines, enabling the analysis of WGS data from different technologies.

## Introduction

Globally tuberculosis is a major cause of mortality with 1.6 million deaths in 2021 (1). It is mainly a disease of the lungs, caused by the bacterium *Mycobacterium tuberculosis* (MTB) (2) and spread from person to person through the air by inhalation of droplets containing MTB (3). TB is most common in developing countries reporting over 95% of cases and deaths (4). MTB strains can become resistant to one or more antibiotics by the acquisition of mutations. Multidrug-resistant tuberculosis (1) is defined as a form of TB that is resistant to first-line anti-TB drugs Isoniazid (5) and Rifampin (6). Patients with MDR-TB are treatable and curable by using second-line drugs, including Fluoroquinolones and Aminoglycosides (7). Second-line treatment requires extensive chemotherapy for at least 2 years which is very expensive and toxic. Additional drug resistance i.e. extensively drug-resistant TB (XDR-TB) is defined as MDR-TB resistant to any or all of the second-line injectable drugs (7).

In 2021 according to WHO, the resistance to all first-line anti-TB drugs (INH, RIF, Pyrazinamide (PZA) and EMB) was 19%, making MDR-TB a public health crisis and a health security threat. Approximately, 450,000 cases were reported by WHO to be resistant to the most effective first-line anti-TB drug RIF, out of which 141,953 people (31%) already had MDR-TB. It is estimated that only 60% of MDR-TB patients are currently successfully treated globally (2).

Attainments of resistance by spontaneous mutation have been estimated as ∼1 in 10^8^ bacilli for RIF, ∼1 in 10^6^ bacilli for INH, streptomycin and EMB. The frequently reported genes that are known to be linked with resistance against INH, RIF, EMB and streptomycin include *rpsl, rrs, rpoB, inh*A, *kat*G, *aph*C, *gid*B and *emb*CAB. In resistant MTB strains, these genes have shown mutation frequency of 70%, ∼10%, 95%, ∼70%, ∼6%, <10% and uncertain respectively(8).

The rate of mutations leading to drug resistance has also been found to be MTB lineage-specific (9). Among 7 lineages, the lineage 2 (for example Beijing lineage family) is highly associated with drug resistance in MTB and has verified higher mutation rates in vitro studies (10)

Next-generation sequencing (1) can provide a detailed sequence of information for multiple drug target genes as well as whole genomes (11). Enhanced coverage of MDR-TB genomes is needed for comprehensive perspective of prediction of drug resistance, thereby impacting patient outcomes. Sequence analysis of MTB genomes has revealed that variation in specific genes that encode drug targets leads to drug resistance in MTB (5, 8). The common mechanism for resistance in MTB is the accumulation of point mutations (SNPs) in the genes encoding drug targets and drug-resistance arises through the selection of mutants during insufficient treatment (1, 12, 13). The genome size of MTB is 4.4 mega base pairs (14) and it comprises of seven lineages. Out of seven, four lineages are predominant in humans which includes lineage 1-4, i.e Indo-Oceanic, East Asian, East African - Indian and Euro-American (5).

Various tools for MTB resistome identification are available, based on detection of resistance-associated alleles. There are two steps required for this; the first step requires the identification of sequence variants (SV) in target genes compared with the MTB reference genome. The second step is the interpretation of the SV in the context of TB resistance. However, categorization of SV associated with MTB resistance is an evolving field, with an increasing number shown to be associated with phenotypic resistance or impacting treatment success. The WHO MTB mutation catalog 2021 is an example of a database which lists mutations with high confidence of prediction for drug resistance, providing updates based on current evidence (3, 15). MTB drug resistance diagnostic tools that link identification of SV with genotypic resistance need to be regularly updated for accurate interpretations. These include, TBProfiler (http:\\tbdr.lshtm.ac.uk\), KVARQ (http://github.com/kvarq/kvarq/), CASTB (http:\\castb.ri.ncgm.go.jp\CASTB\), Mykrobe Predictor (https:\\github.com\iqbal-lab\Mykrobe-predictor\), and Phy-ResSE (www.phyresse.org). Here we present a pilot study to evaluate the accuracy of the MTB-VCF pipeline to identify SVs in gene targets. For this we selected MTB genomes from MDR-TB strains (n=200) and compared the variant calling using MTB-VCF pipeline with that by different drug resistance calling bioinformatics tools; TBProfiler, KVARQ, CASTB, Mykrobe predictor and Phy-ResSE.

## Methods

We downloaded genomes of 200 MDR-TB clinical isolates obtained from ReSeq-TB project (https://platform.reseqtb.org/). The isolates were identified from ReSeq-TB based on the DST profile which was resistant to RIF and INH (Supplementary table 1).

### MTB-VCF tool

An automated variant calling pipeline termed as MTB-VCF was developed to analyze targeted MTB resistome (collection of antibiotics resistance genes) (Figure 1). The targeted variant calling pipeline script is available on GitHub platform https://github.com/safinaARK/MTB-VCF. The pipeline is built on Python v.3.6 snippet and can be run on multiple raw reads (fastq) files using a single command-line on Linux platform. To rapidly characterize mutations from whole genome sequence raw files (fastq format), we map raw sequences to the H37Rv reference genome (NC_000962.3) and variants were called on the 10 target genes (*rpsl, rrs, rpo*B, *inh*A, *kat*G, *aph*C, *emb*CAB, and *gid*B).

**Figure 1:**
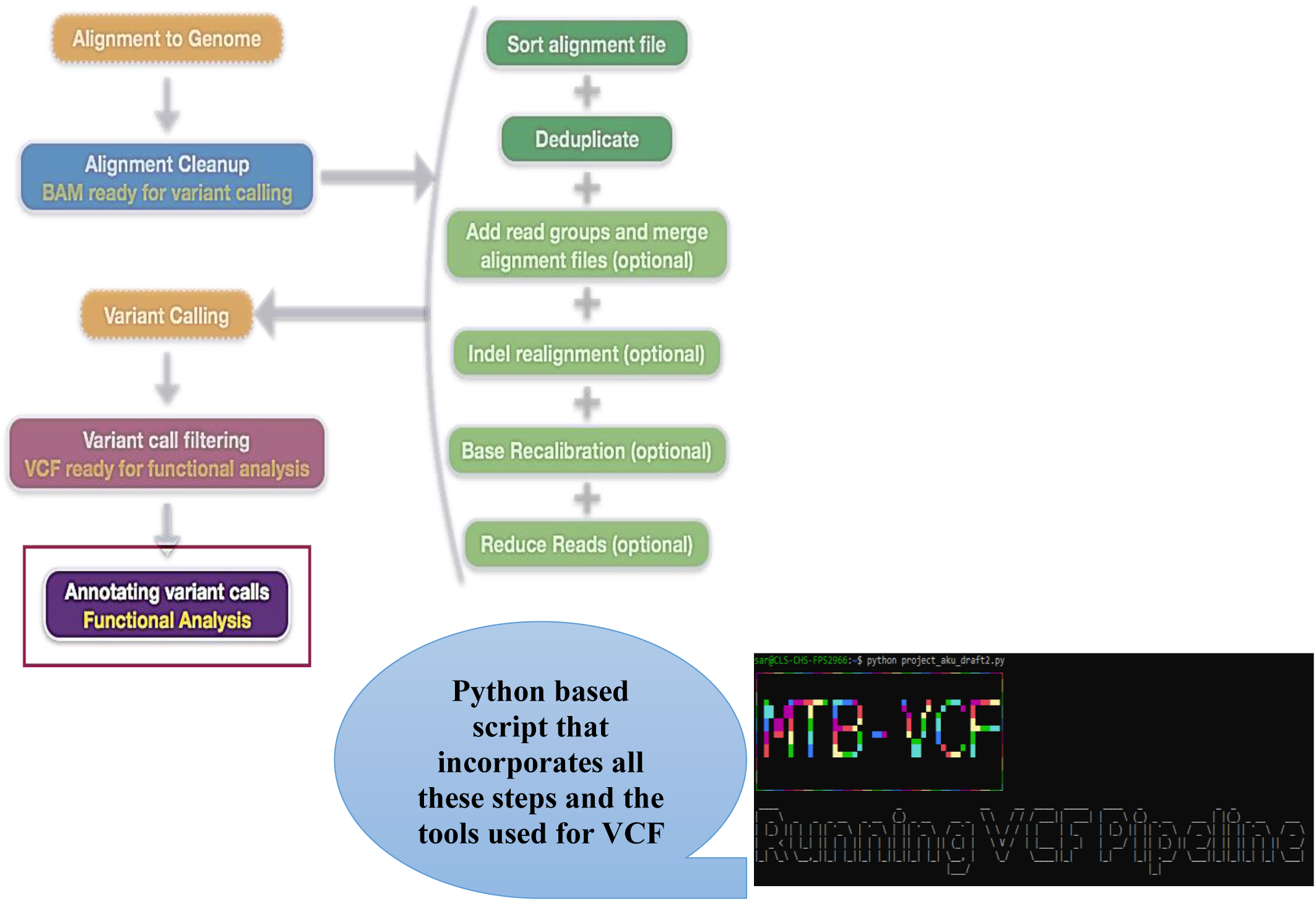
Simplified flow Chart for Automated Targeted Variant Calling (MTB-VCF)

### Comparison of bioinformatics pipelines

We ran the 200 MDR-TB genome datasets on six different tools TB-Profiler v-4.2.0 (Phelan *et. al*., 2019), KvarQ v-0.12.2 (Steiner *et. al*., 2014), CASTB (http://castb.ri.ncgm.go.jp/CASTB/), Mykrobe Predictor TB (https://www.mykrobe.com/), and PhyRes-SE (https://www.phyresse.org/) (Supplementary Figure 1). The SV were called in the 10 genes of interest based on their genomic coordinates, listed in Supplementary Table 1. The genes that are involved in each tool and the SV obtained through the analysis of each tool are listed in Table 1 and Supplementary Table 3.

## Results

### MTB-VCF pipeline evaluation

We used an *in silico* approach using the MTB-VCF pipeline to investigate SV in MTB target genes using a dataset of 200 clinical isolates with an MDR phenotype. MDR-TB isolates were selected from ReSeq-TB based on their phenotypic profile, specifically resistance to INH and RIF, and had a range of DST to EM, SM and Pyrazinamide (PZA) (Supplementary Table 2).

Genomic analysis of the 200 MDR-TB isolates revealed a total of 868 sequence variants, including single nucleotide polymorphisms (SNPs) and insertion/deletion (15) events. Among these variants, 841 were nonsynonymous SNPs (nsSNPs), while 27 were synonymous SNPs (sSNPs) (Table 1). Within the nsSNP category, there were 691 missense SNPs (msSNPs), 8 frameshift SNPs (fsSNPs), 25 stop gain SNPs (sgSNPs), and 53 InDel events (Table 1). For the SV in each target genes, we compared it against those present in the WHO mutation catalog 2021 (3, 15), which were associated with drug resistance. Of the total, 684 SV were the resistance conferring mutation as reported previously.

Notably, the *kat*G gene exhibited the highest number of sequence variants (n=298), followed by *emb*CAB (n=238), *rpo*B (n=163), *rrs* (n=47), *gid*B (n=44), *inh*A (n=33), *ahp*C (n=25), and *rps*L (n=20) (Table 1).

The resistance genotype identified was matched with the phenotypic genotype identified in ReSeqTB (Supplementary Table 2). The sensitivity of resistance identification in the 200 MDR-TB isolates was found to be 99.5% (Table 3).

### Comparison of MTB-VCF with available tools for identification of MTB resistance genotypes

In order to assess the performance of the MTB-VCF pipeline, the number of SVs identified by each tool, including TB-Profiler, KvarQ, CASTB, Mykrobe Predictor, and PhyRes-SE, were assessed and compared with the results obtained from the MTB-VCF pipeline. TB-Profiler identified a total of 653 SVs in 10 genes, while KvarQ identified 89 SVs in 10 genes, CASTB yielded 107 SVs in 5 genes, Mykrobe Predictor identified 62 SVs in 6 genes, and PhyRes-SE identified 110 SVs in 10 genes (Table 2).

The sensitivity and specificity of aforementioned tools in predicting resistance, compared to phenotypic drug susceptibility testing (DST), exhibited considerable variation across different tools and four first-line anti-tuberculosis drug agents (Table 3). The DST profile served as the gold standard for calculating the sensitivity and specificity of each tool for each of the four drugs. Our analysis revealed that the MTB-VCF tool demonstrated higher specificity (99.5%, 95% CI: 98.8-99.9) and sensitivity (97.75%, 95% CI: 96.4-98.9) in calling variants compared to other tools such as TB-Profiler (sensitivity=86%, specificity=95.5%), KvarQ (sensitivity=72.5%, specificity=86.5%), CASTB (sensitivity=79%, specificity=35.75%), Mykrobe Predictor (sensitivity=73.25%, specificity=94.5%), and PhyRes-SE (sensitivity=78.5%, specificity=94.25%). The higher specificity and sensitivity of the MTB-VCF tool were statistically significant, indicating its superior performance in variant calling.

## Discussion

MTB-VCF was implemented as a Python-based command line tool, providing a user-friendly interface for streamlined batch processing of MDR-TB genomes. Raw read (FastQ) files serve as the input for MTB-VCF, identifying SV in the gene of interest as compared reference MTB genome H37Rv. Comparison of SV called by MTB-VCF tool with other bioinformatics pipelines showed a high sensitivity and specificity of variant calling.

We used the tool to identify variants in MDR-TB genomes which were subsequently verified by other bioinformatics pipelines. Several tools for MTB genome sequence analysis and resistance prediction are already available. We compared the results of the MTB-VCF tool with data from five such tools. Most of these tools examine the presence of specific resistance genes or detect SNP and indel mutations associated with drug resistance, thereby predicting the resistance genotypes. TBProfiler is a web-based tool that can also be used via the command line interface. Phy-ResSE and CASTB are exclusively web-based tools. On the other hand, Mykrobe Predictor and KVARQ are software tools that have a Graphical User Interface (16). The results of our analysis clearly demonstrate the superior performance of the MTB-VCF tool in accurately predicting resistance, making it particularly suitable for haploid organisms. The high specificity (99.5%) and sensitivity (97.75%) exhibited by the MTB-VCF tool surpassed those of other evaluated tools such as TB-Profiler, KvarQ, CASTB, Mykrobe Predictor, and PhyRes-SE. These findings highlight the remarkable accuracy of the MTB-VCF tool in variant calling, specifically in the context of drug resistance prediction.

Moreover, our study revealed that the MTB-VCF pipeline exhibited enhanced sensitivity and accuracy in detecting a greater number of SVs compared to the other tools assessed. This comprehensive variant calling approach enables a more in-depth analysis of genomic variations, providing valuable insights into the underlying mechanisms of MDR-TB isolates.

The implementation of cross-platform aware variant calling has been successfully established through the utilization of a genomic VCF workflow. The limitation of the tool is that it cannot be used for resistance prediction as it simply calls variants based on multiple cross platform VCF calling tools incorporated in the pipeline against the MTB reference genome. Another limitation of the MTB-VCF pipeline is that variants identified by it would need to be checked against a mutation database such as the WHO Mutation catalog 2021 (3, 15) to identify its resistance genotype. However, an advantage of the MTB-VCF tool is that it may be applied to any set of genes and can identify various elements, including SNPs, mobile elements, repeated regions, and the MTB-resistome. For example, we have used it to identify variants in efflux pump genes of MTB (17). It also provides flexibility that is lost in predictive platforms that are specifically for a selected list of curated variants. The MTB-VCF platform is highly adaptable, utilizing the genome coordinates of target genes of interest to accurately identify SV without any bias. The variable nature of the MTB-VCF tool enables researchers to investigate and interpret genetic variations in gene targets across a wide range of interests, providing valuable insights into genomic diversity and its implications. This highlights the efficacy and versatility of the MTB-VCF tool in variant calling, emphasizing its potential as a valuable resource for accurate and comprehensive analysis of drug resistance in tuberculosis and other infectious diseases. By accurately identifying and characterizing variants, including SNPs, Indels, and structural variants, the MTB-VCF pipeline enables a more comprehensive understanding of MTB drug resistance mechanisms.

An important distinguishing factor between the MTB-VCF pipeline and the other tools evaluated is the ease with which it can be updated for resistance genes or applied to other pathogens. While the mentioned tools rely on pre-assembled mutation catalogues, the MTB-VCF pipeline has the capability to comprehensively scan all types of polymorphisms (variants, Indels, and structural variants) in the whole genome of any pathogen. Additionally, it generates a variants file using haploid genotypes, further enhancing its versatility and applicability. It is fully customized to predict and annotate variants of targeted genes. It can be modified for customized gene list. MTB-VCF takes a one-line command that can run multiple FASTQ files and generates multi sample VCF files.

In conclusion, the MTB-VCF pipeline can accurately predict drug resistance through calling SV in genes of interest. Its comprehensive variant calling approach makes it a valuable tool for advancing our understanding of MTB drug resistance mechanisms. Its ease of update and adaptability further contribute to its potential as a robust tool in the field of variant calling and genomic analysis. Thus, the MTB-VCF pipeline is a powerful tool for accurate variant calling with potential for advancing our understanding of MTB drug resistance mechanisms.

## Supporting information

Inarticles tables

## Author contributions

SAR wrote the pipeline program, conducted the study, and wrote the first draft. ZH refined the pipeline, guided the study, and written & edited the manuscript. AK analyzed the data and edited the manuscript. SS edited the manuscript. RH obtained the funding and supervised the study. All the authors read and edited the manuscript.

## List of Tables

**Table 1.** Category of Sequence Variants (SV) called by MTB-VCF in each target gene

**Table 2.** Comparison of sequence variants (SV) identified in the MDR-TB dataset (n=200) in tools

**Table 3.** Comparison of the MTB-VCF pipeline with different WGS variant prediction tools based on phenotypic DST to first-line drugs of MDR-TB strains to determine the sensitivity and specificity of the tools

## List of Supplementary Tables

**Supplementary Table 1**. Genes associated with drug resistance in MTB

**Supplementary Table 2**. DST profile of MDR-TB isolates obtained from ReSeq-TB database

**Supplementary Table 3**. Overview of the antibiotics and corresponding resistance genes analyzed by the comparative tools

**Supplementary Table 4**. Sequence Variants associated with drug resistance called by MTB-VCF tool

