## Supplementary material for "A Versatile Tool for Precise Variant Calling in Mycobacterium tuberculosis Genetic Polymorphisms": Inarticles tables

**List of Tables**

**Table 1. Category of Sequence Variants (SV) called by MTB-VCF in each target gene**

| All Sequence Variants identified | | | | | | | | | |
| --- | --- | --- | --- | --- | --- | --- | --- | --- | --- |
| Variants Effect | **rpsL** | **rrs** | **rpoB** | **inhA** | **katG** | **ahpC** | **embCAB** | **gidB** | **Total number of SV** |
| Synonymous | × | × | 7 | 1 | 1 | 1 | 13 | 4 | 27 |
| Missense | 20 | 43 | 133 | 14 | 238 | 8 | 213 | 22 | 691 |
| Frameshift | × | × | × | × | 1 | × | × | 7 | 8 |
| Stop gain | × | × | 3 | × | 20 | × | × | 2 | 25 |
| Intragenic | × | 4 | × | × | × | × | × | × | 4 |
| Upstream | × | × | × | 16 | 10 | 16 | 12 | 5 | 59 |
| Disruptive inframe insertion | × | × | × | 2 | 8 | × | × | 4 | 14 |
| Conservative inframe insertion | × | × | 4 | × | 6 | × | × | × | 10 |
| Conservative inframe deletion | × | × | 15 | × | 14 | × | × | × | 29 |
| Downstream | × | × | 1 | × | × | × | × | × | 1 |
| Total number of SV per gene | 20 | 47 | 163 | 33 | 298 | 25 | 238 | 44 | 868 |

The table depicts 868 SV identified in 200 the MDR-TB genomes analysed

**Table 2. Comparison of sequence variants (SV) identified in the MDR-TB dataset (n=200) in tools**

| Anti-TB drug | genes | MTB-VCF | TB-Profiler | KvarQ | CASTB | Mykrobe Predictor | PhyRes-SE |
| --- | --- | --- | --- | --- | --- | --- | --- |
|  |  | **Total Variants** | | | | | |
| Rifampicin (RIF) | *rpo*B | 163 | 126 | 71 | 51 | 74 | 32 |
| Ethambutol (EMB) | *emb*ABC | 238 | 177 | 5 | 25 | 9 | 33 |
| Isoniazid INH) | *kat*G | 298 | 267 | 6 | 28 | 9 | 13 |
|  | *inh*A | 33 | 26 | 3 | 2 | 6 | 8 |
|  | *aph*C | 25 | 21 | 0 | 0 | 0 | 3 |
| Streptomycin (STR) | *rps*L | 20 | 19 | 4 | 1 | 2 | 3 |
|  | *gid*B | 44 | 2 | 0 | 0 | 0 | 8 |
|  | *rrs* | 47 | 15 | 15 | 0 | 18 | 10 |
| Total SVs |  | 868 | 653 | 104 | 107 | 118 | 110 |

The table lists the number of SV called by each tool across the genes in the 200 MDR-TB genomes analysed.

**Table 3. Comparison of the MTB-VCF pipeline with different WGS variant prediction tools based on phenotypic DST to first-line drugs of MDR-TB strains to determine the sensitivity and specificity of the tools**

| **Anti-TB drug agents** | **DST profile (n=200)** | | **MTB-VCF** | | **TB-Profiler** | | **KvarQ** | | **CASTB** | | **Mykrobe Predictor** | | **PhyRes-SE** | |
| --- | --- | --- | --- | --- | --- | --- | --- | --- | --- | --- | --- | --- | --- | --- |
|  | **#R** | **#S** | **Sens** | **Spec** | **Sens** | **Spec** | **Sens** | **Spec** | **Sens** | **Spec** | **Sens** | **Spec** | **Sens** | **Spec** |
| **INH** | 200 | 0 | 99 | 98 | 93 | 84 | 86 | 95 | 89 | 35 | 89 | 80 | 80 | 98 |
| **RIF** | 200 | 0 | 100 | 98 | 100 | 99 | 94 | 98 | 74 | 45 | 100 | 99 | 90 | 99 |
| **STR** | 164 | 36 | 100 | 95 | 57 | 100 | 57 | 75 | 80 | 38 | 57 | 100 | 50 | 80 |
| **EMB** | 108 | 92 | 99 | 100 | 94 | 99 | 53 | 78 | 73 | 25 | 47 | 99 | 94 | 100 |
| **Overall Sens and Spec** | | | 99.5% | 97.75% | 86% | 95.5% | 72.5% | 86.5% | 79% | 35.75% | 73.25% | 94.5% | 78.5% | 94.25% |

The table classifies the association between the Resistant phenotype and genotype of the 200 MDR TB genomes analyzed (Supplementary Table 2). The resistance prediction was based on the occurrence of resistance-associated SVs as listed in Supplementary Tables 3 and 4.
