## Supplementary Files for "A Versatile Tool for Precise Variant Calling in Mycobacterium tuberculosis Genetic Polymorphisms"

**Supplementary Tables**

**Supplementary Table 1. Genes associated with drug resistance in MTB**

| **Anti-TB Drug** | **Mode of Action** | **Gene** | **MTB genome coordinates** | **Strands** | **Locus Tag** |
| --- | --- | --- | --- | --- | --- |
| Rifampicin | Inhibits biosynthesis cell wall arabinogalactan | rpoB | 759807-763325 | + | Rv0667 |
| Streptomycin | Inhibits protein synthesis by binding to the small subunit bacterial ribosome | rpsL | 780560-782934 | + | Rv0682 |
|  |  | Rrs | 1471846-1473382 | + | MTB000019 |
|  |  | gidB | 4406527-4409201 | - | Rv3919c |
|  |  | KatG | 2152890-2157112 | - | Rv1908c |
| Isoniazid | Inhibits synthesis of cell wall mycolic acid | inhA, | 1673202-1676011 | + | Rv1484 |
|  |  | ahpC | 2726110-2726780 | + | Rv2428 |
| Ethambutol | Inhibits synthesis of proteins | emb operon (embC, embA, embB) | 4240400-4249810 | + | Rv3793, Rv3794, Rv3795, |

**Supplementary Table 2. DST profile of MDR-TB isolates obtained from ReSeqTB database**

|  |  | DST Phenotype | |  |
| --- | --- | --- | --- | --- |
| **ReSeqTB ID** | **RIF** | INH | **ETH** | **STR** |
| ERR038736 | R | R | R | S |
| ERR038737 | R | R | S | S |
| ERR038738 | R | R | R | R |
| ERR038739 | R | R | R | R |
| ERR038740 | R | R | R | S |
| ERR038742 | R | R | R | R |
| ERR038743 | R | R | R | S |
| ERR038744 | R | R | R | R |
| ERR038745 | R | R | R | S |
| ERR038746 | R | R | R | R |
| ERR038747 | R | R | R | R |
| ERR038748 | R | R | R | S |
| ERR038749 | R | R | R | R |
| ERR038750 | R | R | R | R |
| ERR038752 | R | R | S | R |
| ERR038753 | R | R | R | S |
| ERR038754 | R | R | R | R |
| ERR038755 | R | R | S | R |
| ERR040115 | R | R | R | R |
| ERR040119 | R | R | R | S |
| ERR040120 | R | R | R | R |
| ERR040121 | R | R | S | S |
| ERR040122 | R | R | S | S |
| ERR040123 | R | R | R | S |
| ERR040124 | R | R | R | R |
| ERR040125 | R | R | R | S |
| ERR040126 | R | R | R | S |
| ERR040127 | R | R | R | R |
| ERR040128 | R | R | R | R |
| ERR040130 | R | R | R | R |
| ERR040131 | R | R | R | S |
| ERR040132 | R | R | S | S |
| ERR040133 | R | R | R | R |
| ERR040134 | R | R | S | R |
| ERR040135 | R | R | R | R |
| ERR040136 | R | R | R | S |
| ERR040137 | R | R | R | R |
| ERR040138 | R | R | S | R |
| ERR040139 | R | R | R | S |
| ERR040141 | R | R | R | R |
| ERR040142 | R | R | R | S |
| ERR046821 | R | R | S | R |
| ERR046855 | R | R | S |  |
| ERR046917 | R | R | R | S |
| ERR046943 | R | R | S | R |
| ERR067577 | R | R | S | R |
| ERR067579 | R | R | S | R |
| ERR067584 | R | R | R | R |
| ERR067588 | R | R | R | R |
| ERR067589 | R | R | R | R |
| ERR067591 | R | R | S | S |
| ERR067592 | R | R | R | R |
| ERR067595 | R | R | R | R |
| ERR067596 | R | R | S | R |
| ERR067597 | R | R | S | R |
| ERR067598 | R | R | S | R |
| ERR067599 | R | R | R | R |
| ERR067600 | R | R | S | R |
| ERR067602 | R | R | R | R |
| ERR067603 | R | R | S | S |
| ERR067605 | R | R | S | R |
| ERR067611 | R | R | R | R |
| ERR067612 | R | R | R | R |
| ERR067615 | R | R | R | R |
| ERR067616 | R | R | S | R |
| ERR067618 | R | R | R | R |
| ERR067623 | R | R | R | R |
| ERR067624 | R | R | S | R |
| ERR067629 | R | R | R | R |
| ERR067651 | R | R | R | R |
| ERR067653 | R | R | S | R |
| ERR067654 | R | R | R | R |
| ERR067655 | R | R | S | R |
| ERR067656 | R | R | S | R |
| ERR067660 | R | R | R | R |
| ERR067665 | R | R | R | R |
| ERR067673 | R | R | S | R |
| ERR067675 | R | R | R | R |
| ERR067678 | R | R | S | R |
| ERR067680 | R | R | R | R |
| ERR067687 | R | R | S | R |
| ERR067688 | R | R | S | R |
| ERR067698 | R | R | R | R |
| ERR067717 | R | R | R | R |
| ERR067724 | R | R | S | R |
| ERR067726 | R | R | R | R |
| ERR067731 | R | R | S | R |
| ERR067733 | R | R | S | R |
| ERR067734 | R | R | S | R |
| ERR067740 | R | R | S | S |
| ERR067756 | R | R | S | R |
| ERR067757 | R | R | R | R |
| ERR108421 | R | R | R | R |
| ERR108425 | R | R | S | R |
| ERR108436 | R | R | R | R |
| ERR108439 | R | R | R | R |
| ERR108443 | R | R | R | R |
| ERR108458 | R | R | S | R |
| ERR108462 | R | R | R | R |
| ERR108467 | R | R | R | R |
| ERR108488 | R | R | S | R |
| ERR108489 | R | R | S | R |
| ERR108493 | R | R | R | R |
| ERR108500 | R | R | S | R |
| ERR108502 | R | R | S | R |
| ERR108510 | R | R | S | S |
| ERR108515 | R | R | S | R |
| ERR117460 | R | R | R | S |
| ERR117468 | R | R | S | R |
| ERR117470 | R | R | R | R |
| ERR133800 | R | R | S | R |
| ERR133801 | R | R | R | R |
| ERR133803 | R | R | S | S |
| ERR133805 | R | R | S | R |
| ERR133806 | R | R | S | R |
| ERR133815 | R | R | S | R |
| ERR133823 | R | R | R | R |
| ERR133824 | R | R | R | S |
| ERR133827 | R | R | R | R |
| ERR133831 | R | R | R | R |
| ERR133834 | R | R | S | R |
| ERR133836 | R | R | S | R |
| ERR133837 | R | R | S | R |
| ERR133839 | R | R | R | R |
| ERR133842 | R | R | R | R |
| ERR133844 | R | R | R | R |
| ERR133849 | R | R | S | R |
| ERR133858 | R | R | R | R |
| ERR133860 | R | R | S | R |
| ERR133866 | R | R | S | R |
| ERR133868 | R | R | S | R |
| ERR133873 | R | R | S | R |
| ERR133878 | R | R | R | R |
| ERR133880 | R | R | S | S |
| ERR133885 | R | R | R | R |
| ERR133886 | R | R | S | R |
| ERR133888 | R | R | R | R |
| ERR133889 | R | R | R | R |
| ERR133896 | R | R | S | R |
| ERR133902 | R | R | S | R |
| ERR133909 | R | R | S | R |
| ERR133914 | R | R | R | R |
| ERR133915 | R | R | S | R |
| ERR133919 | R | R | R | R |
| ERR133921 | R | R | R | R |
| ERR133922 | R | R | S | R |
| ERR133935 | R | R | R | R |
| ERR133944 | R | R | S | R |
| ERR133953 | R | R | R | R |
| ERR133954 | R | R | R | R |
| ERR133959 | R | R | S | R |
| ERR133962 | R | R | R | R |
| ERR133966 | R | R | S | R |
| ERR133982 | R | R | S | R |
| ERR133984 | R | R | R | R |
| ERR137192 | R | R | R | R |
| ERR137209 | R | R | S | R |
| ERR137210 | R | R | S | S |
| ERR137215 | R | R | S | S |
| ERR137218 | R | R | R | R |
| ERR137222 | R | R | S | R |
| ERR137225 | R | R | S | S |
| ERR137246 | R | R | R | R |
| ERR137247 | R | R | R | R |
| ERR137260 | R | R | S | R |
| ERR137264 | R | R | S | R |
| ERR137269 | R | R | S | R |
| ERR137274 | R | R | S | R |
| ERR137275 | R | R | R | R |
| ERR137277 | R | R | R | R |
| ERR137281 | R | R | S | R |
| ERR137283 | R | R | S | R |
| ERR144548 | R | R | S | R |
| ERR144549 | R | R | S | R |
| ERR144556 | R | R | S | R |
| ERR144560 | R | R | R | R |
| ERR144561 | R | R | S | R |
| ERR144562 | R | R | S | R |
| ERR144563 | R | R | R | R |
| ERR144570 | R | R | S | R |
| ERR144571 | R | R | R | R |
| ERR144574 | R | R | R | R |
| ERR144575 | R | R | S | R |
| ERR144581 | R | R | S | R |
| ERR144590 | R | R | S | R |
| ERR144597 | R | R | S | R |
| ERR144607 | R | R | R | R |
| ERR144619 | R | R | R | S |
| ERR144625 | R | R | S | S |
| ERR144628 | R | R | R | R |
| ERR144634 | R | R | R | R |
| ERR144635 | R | R | R | R |
| ERR158570 | R | R | S | S |
| ERR158576 | R | R | R | R |
| ERR158581 | R | R | R | R |
| ERR158586 | R | R | R | R |
| ERR158592 | R | R | S | S |
| ERR144606 | R | R | R | R |
| ERR158593 | R | R | R | R |
| ERR228019 | R | R | R | S |

Legend – "R" indicates resistance to anti-TB drug agent, while "S" indicates sensitivity.

**Supplementary Table 3. Overview of the antibiotics and corresponding resistance genes analyzed by the comparative tools.**

| **Tool** | **INH** | | | **RIF** | **EMB** | | | **SM** | | |
| --- | --- | --- | --- | --- | --- | --- | --- | --- | --- | --- |
|  | ***ahp*C** | ***inh*A** | ***kat*G** | ***rpo*B** | ***emb*A** | ***emb*B** | ***emb*C** | ***gid*B** | ***rps*L** | ***rss*** |
| CASTB | × | ✓ | ✓ | ✓ | ? | ✓ | ? | ? | ✓ | ? |
| KvarQ | × | ✓ | ✓ | ✓ | × | ✓ | × | × | ✓ | ✓ |
| Mykrobe Predictor | × | ✓ | ✓ | ✓ | × | ✓ | × | × | ✓ | ✓ |
| Phy-ResSE | ✓ | ✓ | ✓ | ✓ | ✓ | ✓ | ✓ | ✓ | ✓ | ✓ |
| TBProfiler | ✓ | ✓ | ✓ | ✓ | ✓ | ✓ | ✓ | ✓ | ✓ | ✓ |
| MTB-VCF | ✓ | ✓ | ✓ | ✓ | ✓ | ✓ | ✓ | ✓ | ✓ | ✓ |

Legend – The Tick (✓) represents the gene that was predicted in the tool whereas, the cross (×) denotes that the genes are not included. The Question marks (?) were used for CASTB where it was unclear whether the genes are interrogated.

**Supplementary Table 4. Sequence Variants associated with drug resistance called by MTB-VCF tool**

| Resistance conferring Variants | | | | | | | | | |
| --- | --- | --- | --- | --- | --- | --- | --- | --- | --- |
| Variants Effect | ***rps*L** | ***rrs*** | ***rpo*B** | ***inh*A** | ***kat*G** | ***ahp*C** | ***emb*CAB** | ***gid*B** | **Total number of SV** |
| Synonymous | × | × | 1 | × | × | × | × | × | 1 |
| Missense | 17 | 43 | 104 | 13 | 222 | 7 | 176 | 7 | 589 |
| Frameshift | × | × | × | × | × | × | × | × | 0 |
| Stop gain | × | × | × | × | 19 | × | × | 1 | 20 |
| Intragenic | × | × | × | × | × | × | × | × | 0 |
| Upstream | × | × | × | 11 | 3 | 15 | 4 | × | 33 |
| Disruptive inframe insertion | × | × | × | × | 4 | × | × | × | 4 |
| Conservative inframe insertion | × | × | 4 | × | 5 | × | × | × | 9 |
| Conservative inframe deletion | × | × | 15 | × | 13 | × | × | × | 28 |
| Downstream | × | × | 0 | × | × | × | × | × | 0 |
| Total number of SV per gene | 17 | 43 | 124 | 24 | 266 | 22 | 180 | 8 | 684 |

Legend - Cross (×) indicates the absence of a specific sequence variation (SV), while the numbers represent the quantity of each SV present in a particular gene.
